# High altitude adaptation mitigates anemia risk associated with diabetes among the Mosuo of Southwest China

**DOI:** 10.1101/406579

**Authors:** M Su, K Wander, MK Shenk, T Blumenfield, H Li, SM Mattison

## Abstract

Human populations native to high altitude regions (≥2500 m) exhibit numerous adaptations to hypoxic stress. On the Tibetan Plateau, these include modifications of the hypoxia inducible factor (HIF) pathway to essentially uncouple erythropoiesis (red blood cell production) and blood hemoglobin (Hb) concentration—which normally increase in response to low oxygen—from hypoxia. Uncoupling of erythropoiesis and hypoxia is also observed among people with diabetes due to damage to kidney tissues. This is hypothesized to result in elevated risk for anemia among diabetics, which increases risk for cardiovascular disease and death. We tested the hypothesis that the independence of erythropoiesis from HIF among high-altitude adapted populations of the Tibetan Plateau may protect against diabetes-associated anemia. We investigated this hypothesis among the Mosuo, a population living in Yunnan Province, China (at ~2800 m altitude) that is undergoing rapid market integration and lifestyle change, with concomitant increase in risk for type 2 diabetes. We found that, although diabetes (glycated hemoglobin, HbA_1c_ ≥6.5%) is associated with anemia (females: Hb<12g/dl; males: Hb<13g/dl) among the Chinese population as a whole (N: 5,606; OR: 1.48; *p*: 0.008), this is not the case among the Mosuo (N: 316; OR: 1.36; *p*: 0.532). Both pathways uncoupling hypoxia from erythropoiesis (diabetic disease and high altitude adaptation) are incompletely understood; their intersection in protecting Mosuo with diabetes from anemia may provide insight into the mechanisms underlying each. Further, these findings point to the importance of understanding how high-altitude adaptations interact with chronic disease processes, as populations like the Mosuo experience rapid market integration.

## INTRODUCTION

Diabetes dramatically increases risk for anemia. There are multiple possible pathways linking diabetes and anemia, including higher rates of inflammation, autoimmunity, and micronutrient deficiency among those with diabetes. However, one pathway unique to diabetes has been repeatedly implicated: failure of the erythropoietin response to hypoxia in the kidney [1-3]. This seems to occur prior to other manifestations of diabetic kidney disease, and is not a similarly early manifestation of non-diabetic renal disease [2, 3]. Here, we assess the extent to which diabetes increases risk for anemia among the Mosuo ethnic group in China, a population living at ~2800m in altitude in the Hengduan Mountains in Yunnan Province [4], where we hypothesize genetic adaptations to high altitude [5, 6] may disrupt diabetes’ unique ability to cause anemia via the erythropoietin pathway.

### High altitude adaptation

Low barometric pressure and low oxygen concentration at altitudes ≥2500 m create hypoxic (low oxygen) stress. Indigenous populations of the Tibetan Plateau have been living at altitudes of up to ~5400 m—under constant hypoxic stress—for thousands of years [5]. Tibetans exhibit a suite of characteristics that appear to fully mitigate this stress: while sojourners to high altitude experience a ~10-20% deficit in maximal oxygen uptake (and thus ability to do work requiring oxygen delivery to their tissues), Tibetans (as well as populations native to the Andean Plateau) at high altitude exhibit no deficit compared to lowland natives at low altitude [5]. This adaptation to high altitude is accomplished through multiple physiological changes from the low-altitude norm.

Tibetans exhibit a suite of adaptations to high altitude. These include higher resting ventilation than lowland natives at low altitude (similar to the elevated respiration rate seen temporarily among sojourners to high altitude) [5, 6]. Vasodilation, attributable to higher circulating nitric oxide (NO), and denser capillary networks among Tibetans allow high rates of blood flow to tissues, which may compensate for lower arterial blood oxygen content [6, 7].

While dramatic increases in blood hemoglobin (Hb) are typical when lowland natives move to high altitude (or are raised at high altitude) and are part of the adaptive suite of populations indigenous to the Andean Plateau, Tibetans exhibit Hb that is only somewhat higher than lowland Hb at low attitude [5, 6]. Tibetans also have lower erythropoietin (EPO), which stimulates red blood cell production, than Andeans [5, 6]. Tibetans have lost some responses to acute hypoxia that can be harmful when activated long term. Their blunted erythropoietic response to hypoxic stress, moderate Hb, and low EPO (compared to sojourners and Andeans) avoids blood hyperviscosity that would come from high rates of erythropoiesis and a large red blood cell population [5, 6]. Hyperviscostiy can decrease blood flow and profusion, and can cause retinopathy, bleeding or hemorrhage, as well as neurological symptoms, including seizures. Tibetans also exhibit no hypoxic pulmonary vasoconstriction—a normal response to local hypoxia in the lung, which redirects blood to better oxygenated areas of the lungs—due to high levels of blood and lung NO [5]. Hypoxic pulmonary vasoconstriction can lead to life-threatening pulmonary hypertension and pulmonary edema.

Multiple studies have identified evidence of positive natural selection in the Tibetan genome, including on genes associated with the hypoxia-inducible factor (HIF) pathway of erythropoiesis. (Other pathways were implicated as well [8-12]; here, we focus on the HIF pathway for its relevance to anemia and diabetes.) HIF pathway genes identified as under selection included PHD2 (prolyl hydroxylase domain protein 2, also known as ELGN1, Egl nine homolog 1), PPARA (peroxisome proliferators-activated receptor α), [8] and HIF2A (HIF-2α, also known as EPAS1, endothelial PAS domain protein 1) [9, 10]. Alleles at high frequency among Tibetans (relative to Han) were inversely associated with Hb [8- 10]. This suggests that, in the presence of physiological changes that alleviate the need for elevated erythropoiesis (such as vasodilation and increased blood flow, resulting from as yet poorly understood genetic and/or developmental pathways), natural selection has favored changes to the HIF pathway that uncoupled erythropoiesis from hypoxia sensing, allowing Tibetans to adapt to high altitude environments without incurring the cost of elevated red blood cell production (*vs*. that normal at lower altitudes) and blood hyperviscosity.

Hypoxia inducible factor (HIF) is regulated primarily, but not exclusively, by oxygen concentrations: The prolyl hydroxylase domain enzymes (PHD1-3) hydroxylate the α subunit of HIF in an oxygen-dependent manner—as long as there is adequate oxygen, the α subunit of HIF is rare. This hydroxylation ceases under hypoxic conditions, allowing HIF-α to stabilize and dimerize with HIF-β to form HIF [6]. HIF can then activate numerous genes. Three types of HIF exist, differing in their α subunit (HIF-α1, HIF-α2, HIF-α3); HIF-α2 regulates erythropoietin (EPO) expression in the kidney, and EPO regulates erythropoiesis in the bone marrow [6]. It makes sense, then, given how central the HIF pathway is to responding to hypoxia, that two elements of this pathway—PHD2 (or ELGN1), which links oxygen conditions to HIF, and HIF2 (or EPAS1), which extends this link to EPO—are involved in Tibetans’ unique adaptations to high altitude. More specifically, it has been hypothesized that the adaptive allele(s) of PHD2 and/or HIF2 involve loss of function mutation(s) that break the links between hypoxia and EPO [6].

One aspect of high altitude adaptation that remains relatively under-researched is the impact of high altitude adaptation on chronic diseases. Particularly, as social change occurs in high altitude populations—e.g., market integration and growing inequality; changes in subsistence, diets, and physical activity—how is risk for chronic disease affected? How do physiological adaptations to high altitude interact with chronic disease epidemiology, etiology, complications, and treatment? Here, we investigate diabetes, and associated risk for anemia, among the Mosuo, close relatives of Tibetans living in the Hengduan Mountains southeast of the Tibetan Plateau, who are undergoing a rapid process of market integration associated with increasing tourism in the area around Lugu Lake in Yunnan Province.

### Anemia among diabetics

Diabetes is a disease that is defined by elevated blood glucose, and caused by impaired production of, or sensitivity to, the hormone insulin, which promotes tissue absorption of glucose. In type 1 diabetes (T1D), insulin production in the pancreas is impaired by autoimmune attack on insulin-producing beta cells. In type 2 diabetes (T2D), cells cease to respond to normal levels of insulin (insulin insensitivity or insulin resistance), leading to, among other manifestations, release of glucose stored in the liver, and decreasing insulin production in the pancreas. High and increasing prevalences of T2D are a global public health concern [13], with particularly dramatic change among populations like the Mosuo, undergoing rapid integration into markets, with attendant changes in diets, nutrition, physical activity, social structure, wealth, and inequality [14, 15].

Diabetes’ (T1D and T2D) effects on the body are numerous and distributed across many organ systems. Common complications of diabetes include cardiovascular disease, nerve damage, blindness, and kidney failure. Although not consistently emphasized in discussions of diabetes complications, anemia is also very common among people with both T1D and T2D. (For example, 32% in [16]; 23% in [17]; 11.6% in [18], using cutpoints lower than those recommended by WHO; and 14% among T1D patients alone in [19]). Moreover, anemia increases risks for severe diabetes complications, such as CVD and death [20].

It is likely that anemia arises from multiple causes among people with diabetes, including autoimmunity, inflammation, and nutritional deficiency. The most salient cause of anemia among those with diabetes, however, may be their impaired ability correct insipient anemia via the homeostatic kidney erythropoietin (EPO) response. Although anemia among people with diabetes is not limited to those with diabetic nephropathy, it is particularly common among this group, and can occur very early in diabetic nephropathy [1, 2].

The HIF hypoxia sensing system in the healthy and normally-functioning kidney maintains oxygen homeostasis by increasing EPO production by peritubular fibroblasts in response to hypoxia and vasoconstriction, which stimulates proliferation and differentiation of erythrocytes in bone marrow [2, 21]. This homeostatic system increases EPO in responses to low Hb, correcting emerging anemia with increased red blood cell production. When the system works properly, a generally inverse relationship between Hb and EPO is observed; when this correction fails, as seems to be the case in anemic diabetics, the inverse relationship between EPO and Hb is not observed, and EPO is inappropriately low among people with anemia, compared to those without.

Kidney damage in diabetes appears to occur primarily via damage to the proximal tubule [3]. In particular, peritubular fibroblasts (which produce EPO) may be damaged by hyperglycemia, increased capillary pressure, or proteinuria, resulting in fibrosis [2]. This fibrosis may uncouple renal EPO production from hypoxia, so that EPO remains inappropriately low (i.e., in the normal range) during anemia [3]. The failure of EPO elevation is observed among anemic diabetics, both with and without diabetic nephropathy [2, 3]. Because this fibrosis is unlikely to result in severe damage, and because anemia and under-production of EPO among those with diabetes often precede other signs of declining renal function, and finally, because EPO production among anemic diabetics is not compromised below normal (non-anemic) levels until kidney disease becomes severe, *sensitivity* to hypoxia, rather than EPO secretory capacity, is likely to be the compromised element of EPO regulation that is responsible for anemia among those with diabetes [1, 3, 21].

### Hypothesis and predictions

If Tibetans’ suite of adaptations to high altitude, including increased respiration and enhanced blood flow (the genetic bases of which are only beginning to be understood) decrease the need for hemoglobin at altitude, alleles of PHD2 and HIF2 that uncouple hypoxia from erythropoiesis could have provided a fitness advantage by decreasing red blood cell production into the normal (low altitude) range, decreasing risk for adverse effects of blood hyperviscosity and elevated hemoglobin. Tibetans’ adaptations to high altitude, and the independence of erythropoiesis from the HIF pathway, may have the additional advantage of protecting those who develop diabetes against anemia.

If this is the case, we predict that the elevated anemia risk normally associated with diabetes will be present in the Chinese population, but not observed in a population with Tibetan-type high-altitude adaptations. Similarly, we predict that diabetes will be associated with lower Hb only among the general Chinese population. (Using Hb as a continuous variable allows us to avoid confusion over the appropriateness of WHO anemia definitions, and particularly correction for altitude, given that only moderate increases in Hb are observed at high altitude among Tibetans.) We test these predictions (Table 1) among the Mosuo, whose recent Tibetan ancestry and rapid market integration bring together high altitude adaptation and high diabetes risk, allowing us to observe how the two interact to affect anemia.

**Table 1.** Hypothesized effect of diabetes on anemia risk and hemoglobin.

|  | <u>Anemia risk</u> | <u>Hemoglobin concentration</u> |
| --- | --- | --- |
| CHNS | + | - |
| Mosuo | None | None |

## MATERIALS & METHODS

### Study setting and design

Associations between diabetes and anemia among the Chinese population as a whole were described with the China Health and Nutrition Survey (CHNS: http://www.cpc.unc.edu/projects/china). Associations between diabetes and anemia among the Mosuo were described as part of a larger survey of the Mosuo, investigating the impact of rapid market integration on wealth, health, and inequality. Rapid market integration is occurring among some Mosuo communities due to increasing tourism around Lugu Lake in Yunnan Province. Communities were selected to include those with high, moderate, and very low levels of participation in the tourism economy.

The demographic survey portion of the Mosuo study was carried out in 2017. A sample of participants in the survey were invited to participate in screening for diabetes and anemia after their initial participation; screening was carried out in the summers of 2017 and 2018. Individual participants were not selected at random; instead, a subset of communities from the larger survey were targeted. Representation of communities with varying degrees of access to market economies was ensured, from those participating in the emerging tourism economy to those that remain isolated by remote locations and limited infrastructure. All adult participants in those communities were invited to provide specimens and be screened for anemia and diabetes, and all of those who volunteered were included in the subsample. All participants provide verbal informed consent.

Aspects of health—particularly, inflammation, diabetes, iron deficiency, and anemia—were described among this sub-sample using biomarkers. Biomarkers for these health outcomes were selected to be measurable, when possible, with field-friendly point-of-care test (POCT) devices, and, when this was not possible, to be measurable in dried blood spot (DBS) specimens. Biomarkers were also selected for interpretability in a population with a high burden of infectious disease (e.g., robustness of the iron nutrition biomarker to changes in inflammation), and for practicality in population-based research (e.g., minimal diurnal variation in all biomarkers, minimal pre-test preparation requirements for participants).

Inflammation was described with C-reactive protein, a stable, commonly-used, and easy to measure pro-inflammatory acute phase reactant. Diabetes was characterized with glycated hemoglobin (HbA_1c_). HbA_1c_ has been validated against blood glucose measures as gold standards [13]; it was selected for use in this project to meet the practical limitations of population-based research, as it does not require research participants to fast prior to participation, and is less variable over short time scales than fasting plasma glucose or oral glucose tolerance tests [22]. Limitations of HbA1c include artificially *low* values for individuals with elevated erythrocyte turnover (e.g., due to hemolytic disease, hemoglobinopathy, or recent acute blood loss), and artificially *high* values for individuals with iron deficiency (with or without anemia) [22, 23]. Iron deficiency was described with soluble transferrin receptor (sTfR), which, unlike many other biomarkers of iron status, is unaffected by inflammation and the acute phase response, enhancing its interpretability in settings with a high infectious disease burden, and is measurable in DBS [24, 25]. Anemia was described with hemoglobin (Hb) [26].

### Anthropometry

At the time of participation in the demographic survey of the Mosuo, each participant’s height was measured with a stadiometer and weight with a digital scale. Waist circumference was measured with a body tape [27].

### Point of care testing

Capillary whole blood was collected via finger stick with a sterile safety lancet. Blood specimens were immediately evaluated for HbA_1c_ with the A1CNOW POCT (PTS Diagnostics) and Hb with a hemoglobinometer (HemoCue 201+).

### Laboratory evaluation of biomarkers

After POCT, additional drops of whole blood were allowed to fall freely onto a filter paper card (Whatman #903) for dried blood spot (DBS) specimens. DBS were allowed to dry for up to 24h and were then frozen. Specimens were transported to the MOE Key Laboratory for Contemporary Anthropology at Fudan University and evaluated for C-reactive protein (CRP; BioCheck BC-1119) and soluble transferrin receptor (sTfR; Ramco TFC-94) using commercially available kits modified for use with DBS specimens: One 1/8 inch disc of DBS specimen was removed with a hole punch, combined with the specimen dilution buffer provided with each kit, and allowed to soak overnight at 4°C. The resulting eluent was assayed without further dilution; dilution was calculated as 1.525 μl serum equivalent [28] per volume of dilution buffer.

### CHNS data access

The CHNS “Biomarker Data” dataset was obtains via download from the study website. These data include biomarkers information from specimens collected in 2009 and associated individual-level information (e.g., age, sex, smoking behavior).

### Data analysis

Diabetes was defined according to WHO standard as HbA_1c_ ≥ 6.5% [13]. This definition captures only those with unrecognized or poorly controlled diabetes, but excludes those with well-controlled diabetes. Among the Mosuo, access to care is limited and few with diabetes are either aware of their condition or being assisted by a medical care provider to control it. As such, we defined diabetes among the Mosuo (and, for comparability, CHNS) only on the basis of HbA_1c_. Anemia was defined according to WHO standards [26] both with (as Hb ≤ 13.3 g/dl for women and Hb ≤ 14.3 g/dl for men) and without (as Hb ≤ 12 g/dl for women and Hb ≤ 13 g/dl for men) adjustment of 1.3 g/dl for elevation of 2500 m. The unadjusted reference may be more appropriate for the Mosuo, as high-altitude adapted

Tibetans have been found to have only slightly elevated Hb compared to those at sea level. Iron deficiency was defined differently in CHNS and the Mosuo datasets: Unlike other biomarkers, sTfR measurement methods can generate tremendously different values [25], and cutpoints are specific to analysis method. sTfR in CHNS was evaluated with nephelometry (Siemens BNP), rather than enzyme immunoassay, as was used among the Mosuo. Thus, iron deficiency was defined as sTfR > 1.76 mg/l in CHNS and sTfR ≥ 8.3 mg/l among the Mosuo. sTfR was also evaluated as a continuous variable, as was CRP. Body mass index was calculated as weight (g)/height (m)^2^, and BMI was evaluated as a continuous variable.

Associations between diabetes and anemia were described with logistic regression using Stata 15 software (StataCorp). Models were constructed separately for the Mosuo and the CHNS datasets. Known anemia risk factors that were also likely to be associated with diabetes or the HbA_1c_ test result were included as control variables: smoking, sex, age, sTfR, CRP, and body mass index (BMI). Province was also included as a control variable in the model of CHNS data; field season (2018 *vs*. 2017) was include as a control variable in the model of Mosuo data. Limited access to health care among the Mosuo, and limited communication in the healthcare setting, made it difficult for participants to accurately describe their current medications and co-morbid conditions; as such, we excluded these additional possible causes of confounding from our analyses.

Linear regression was used to describe associations between Hb and diabetes (with the same control variables as logistic regressions). Modeling Hb as a continuous variable avoids uncertainty regarding the appropriate cut-points for anemia for populations with Tibetan-type high-altitude adaptations, but is less interpretable as regards the clinically meaningful outcome of anemia.

## RESULTS

Complete CHNS information was available for 5,606 individuals (Table 2). The prevalence of diabetes in CHNS was 7.47%; anemia was also common (14.88%).

**Table 2.** China Health and Nutrition Survey sample characteristics

|  | Number | % |
| --- | --- | --- |
| Anemia (Hb < 12 for females and Hb < 13 for males) | 834 | 14.88 |
| Diabetes (HbA<sub>1c</sub> ≥ 6.5%) | 419 | 7.47 |
| Iron deficiency (sTfR ≥ 1.76 mg/L) | 1,050 | 18.70 |
| Current smoking | 2,146 | 38.28 |
| Female sex | 2,784 | 49.66 |
|  | Median | Range |
| Age (years) | 51.52 | 18.03, 94.45 |
| C-reactive protein (mg/L) | 1 | 0, 500 |
| Soluble transferrin receptor (mg/L) | 1.33 | 0.13, 14.20 |
| Body mass index | 21.77 | 12.33, 37.88 |

Complete information was available for 316 Mosuo (Table 3). Diabetes was common among the Mosuo (14.56%). Anemia was common as well, whether the definition was adjusted (29.75%) or unadjusted (10.76%) for altitude. 11.39% of participating Mosuo were iron deficient. Females were over-represented in this sample.

**Table 3.** Mosuo sample characteristics

|  | Number | % |
| --- | --- | --- |
| Anemia |  |  |
| Adjusted for altitude (Hb < 13.3 for females and Hb < 14.3 for males) | 94 | 29.75 |
| Unadjusted for altitude (Hb < 12 for females and Hb < 13 for males) | 34 | 10.76 |
| Diabetes (HbA<sub>1c</sub> ≥ 6.5%) | 46 | 14.56 |
| Iron deficiency (sTfR ≥ 8.3 mg/L) | 36 | 11.39 |
| Iron deficiency anemia (adjusted for altitude) | 9 | 2.85 |
| Iron deficiency anemia (unadjusted for altitude) | 3 | 0.95 |
| Current smoking | 73 | 23.10 |
| Female sex | 215 | 68.04 |
|  | Median | Range |
| Age (years) | 47 | 14-84 |
| C-reactive protein (mg/L) | 0.79 | 0.05, 52.64 |
| Soluble transferrin receptor (sTfR) | 6.17 | 2.93, 16.78 |
| Body mass index | 23.93 | 15.68, 33.76 |

As predicted, diabetes was positively associated with anemia in the CHNS data (Table 4). Controlling for smoking, sex, age, province, sTfR, CRP, and BMI, diabetes (HbA_1c_ ≥ 6.5%) was associated with 50% higher odds of anemia (OR: 1.48, *p*: 0.008).

**Table 4:** Logistic regression model of risk factors for **anemia (Hb < 12 for females and Hb < 13 for males)** in CHNS (N = 5,606)

|  | OR | 95% CI | P |
| --- | --- | --- | --- |
| Diabetes (HbA<sub>1c</sub> ≥ 6.5%) | 1.48 | 1.11, 1.99 | 0.008 |
| Smoking (current) | 1.14 | 0.96, 1.35 | 0.127 |
| Female sex | 2.24 | 1.89, 2.65 | 0.000 |
| Age | 1.03 | 1.02, 1.04 | 0.000 |
| Province (Reference: Liaoning) |  |  |  |
| Heilongjiang | 1.44 | 0.93, 2.25 | 0.106 |
| Jiangsu | 2.08 | 1.42, 3.05 | 0.000 |
| Shandong | 1.76 | 1.16, 2.66 | 0.008 |
| Henan | 1.79 | 1.17, 2.75 | 0.008 |
| Hubei | 3.59 | 2.44, 5.28 | 0.000 |
| Hunan | 4.38 | 3.00, 6.39 | 0.000 |
| Guangxi | 1.91 | 1.29, 2.83 | 0.001 |
| Guizhou | 1.21 | 0.78, 1.86 | 0.397 |
| Soluble transferrin receptor (sTfR) | 3.00 | 2.64, 3.40 | 0.000 |
| C-reactive protein (CRP) | 1.00 | 1.00, 1.01 | 0.279 |
| Body mass index (BMI) | 0.92 | 0.89, 0.95 | 0.000 |

Among the Mosuo, diabetes was *not* positively associated with anemia. Controlling for smoking, sex, age, CRP, sTfR, BMI, and field season, diabetes was unassociated with anemia (whether or not anemia definitions were adjusted for altitude; Tables 5, 6).

**Table 5:** Logistic regression model of risk factors for **anemia (Hb < 13.3 for females and Hb < 14.3 for males)** among Mosuo (N = 316)

|  | OR | 95% CI | P |
| --- | --- | --- | --- |
| Diabetes (HbA<sub>1c</sub> ≥ 6.5%) | 0.56 | 0.25, 1.23 | 0.149 |
| Smoking (current) | 0.56 | 0.22, 1.45 | 0.233 |
| Female sex | 4.92 | 2.08, 11.61 | 0.000 |
| Age in years | 0.99 | 0.98, 1.01 | 0.539 |
| Soluble transferrin receptor (sTfR) | 0.95 | 0.83, 1.08 | 0.435 |
| C-reactive protein (CRP) | 0.99 | 0.91, 1.07 | 0.764 |
| Body mass index (BMI) | 0.97 | 0.90, 1.05 | 0.484 |
| Field season (2018) | 0.42 | 0.24, 0.78 | 0.003 |

**Table 6:**
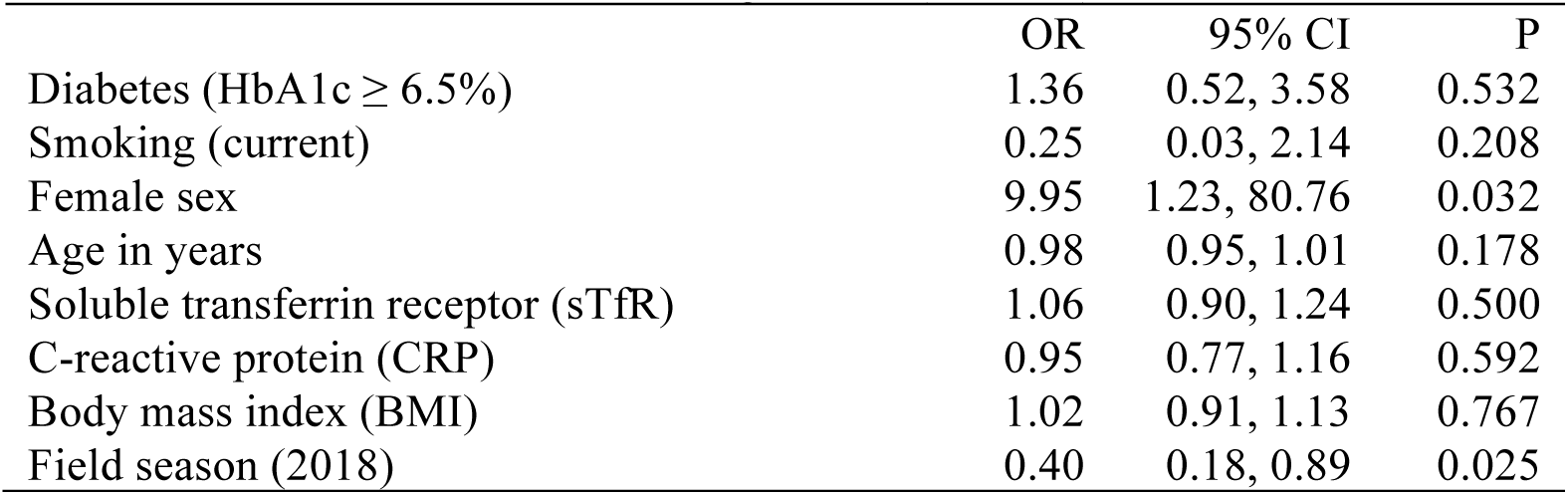
Logistic regression model of risk factors for **anemia (Hb < 12 for females and Hb < 13 for males)** among Mosuo (N = 316)

|  | OR | 95% CI | P |
| --- | --- | --- | --- |
| Diabetes (HbA1c ≥ 6.5%) | 1.36 | 0.52, 3.58 | 0.532 |
| Smoking (current) | 0.25 | 0.03, 2.14 | 0.208 |
| Female sex | 9.95 | 1.23, 80.76 | 0.032 |
| Age in years | 0.98 | 0.95, 1.01 | 0.178 |
| Soluble transferrin receptor (sTfR) | 1.06 | 0.90, 1.24 | 0.500 |
| C-reactive protein (CRP) | 0.95 | 0.77, 1.16 | 0.592 |
| Body mass index (BMI) | 1.02 | 0.91, 1.13 | 0.767 |
| Field season (2018) | 0.40 | 0.18, 0.89 | 0.025 |

Linear regressions show no association between diabetes and Hb in either dataset (Tables 7, 8).

**Table 7:** Linear regression model of predictors of **hemoglobin (g/dl)** for CHNS

| | $\beta$ | 95% CI | P |
| --- | --- | --- | --- |
| Diabetes (HbA1c ≥ 6.5%) | 0.05 | -0.13, 0.23 | 0.582 |
| Smoking (current) | -0.07 | -0.17, 0.02 | 0.131 |
| Female sex | -1.90 | -1.99, -1.81 | 0.000 |
| Age in years | -0.02 | -0.02, -0.02 | 0.000 |
| Province (Reference: Liaoning) |  |  |  |
| Heilongjiang | -0.50 | -0.71, -0.29 | 0.000 |
| Jiangsu | -0.00 | -0.19, 0.19 | 0.974 |
| Shandong | -0.13 | -0.33, 0.07 | 0.208 |
| Henan | -0.44 | -0.65, -0.23 | 0.000 |
| Hubei | -0.95 | -1.15, -0.74 | 0.000 |
| Hunan | -0.98 | -1.18, -0.78 | 0.000 |
| Guangxi | -0.50 | -0.70, -0.30 | 0.000 |
| Guizhou | -0.32 | -0.53, -0.11 | 0.003 |
| Soluble transferrin receptor (sTfR) | -0.65 | -0.71, -0.59 | 0.000 |
| C-reactive protein (CRP) | -0.00 | -0.01, 0.00 | 0.931 |
| Body mass index (BMI) | 0.05 | 0.03, 0.06 | 0.000 |

**Table 8:** Linear regression model of predictors of **hemoglobin (g/dl)** for Mosuo

| | $\beta$ | 95% CI | P |
| --- | --- | --- | --- |
| Diabetes (HbA1c ≥ 6.5%) | -0.09 | -0.44, 0.62 | 0.741 |
| Smoking (current) | 0.55 | -0.00, 1.10 | 0.051 |
| Female sex | -2.37 | -2.88, -1.87 | 0.000 |
| Age in years | 0.01 | -0.01, 0.02 | 0.249 |
| Soluble transferrin receptor (sTfR) | 0.05 | -0.04, 0.15 | 0.247 |
| C-reactive protein (CRP) | -0.02 | -0.06, 0.03 | 0.402 |
| Body mass index (BMI) | 0.04 | -0.01, 0.10 | 0.095 |
| Field season (2018) | 0.52 | 0.13, 0.91 | 0.009 |

## DISCUSSION

We found that, although anemia is more common among those with diabetes in the Chinese population in general, this is not the case among the Mosuo, a group living at ~2800 m in altitude in the Hengduan Mountains southeast of the Tibetan Plateau. The Mosuo are undergoing rapid market integration, with concomitant increasing risk for diabetes and other chronic diseases. We postulate that this protection from diabetes-associated anemia arises from adaptations to high altitude that enhance oxygen delivery to tissues in the absence of dramatic increases in red blood cell production, and uncouple erythropoiesis regulation from hypoxia. In short, for most lowland people with diabetes, diabetic kidney damage compromises the kidneys’ ability to sense (and so respond to) hypoxia, disrupting a system that normally works to prevent anemia. For the Mosuo, however, high altitude adaptations may have already uncoupled erythropoiesis regulation from hypoxia, protecting Mosuo with diabetes against this manifestation of disease.

In this respect, physiological mechanisms that uncouple erythropoiesis from hypoxia sensing among the Mosuo may provide them some protection in an increasingly diabetogenic environment: they are protected against anemia as an early complication of diabetes (and possibly the increased risk for cardiovascular disease and death associated with anemia among diabetics).

### Limitations

The Mosuo sample was not selected at random from among the entire Mosuo population, and so its generalizability is limited. Those who chose to participate may have been motivated by symptoms of disease, and so we may have included those with anemia or diabetes at a higher rate than those without. Thus, the fairly high rates of diabetes and anemia we observed must be interpreted with great caution. However, it seems less likely that bias introduced by higher rates of participation by those with symptoms of diabetes or anemia could have affected our main finding (no association between diabetes and anemia) by obscuring a real, positive association between diabetes and anemia.

We selected the HbA_1c_ biomarker to identify diabetes for multiple practical reasons: it can be measured at the point of care and without any preparation by the participant; contrary to screenings that rely on plasma glucose measurement, it exhibits no short-term intra-individual variation (e.g., with time of day, recent food ingestion, exercise, or acute stress—all reflecting the dynamic process of glucose homeostasis); and, it is more strongly associated with subsequent development of complications among diabetes patients than fasting plasma glucose [22]. However, defining diabetes by HbA_1c_ introduces two possible sources of bias into our analyses, both arising from the fact that HbA_1c_ is only interpretable when erythropoiesis and erythrocyte lifespan are normal.

First, it is possible that some aspect of adaptation to high altitude has altered erythropoiesis or erythrocyte lifespan among the Mosuo in a way that increases HbA_1c_. We think this is unlikely, as many lines of evidence summarized above suggest that elevated erythropoiesis is not part of the Tibetan pattern of adaptation to high altitude [5, 6]. However, since both erythropoiesis and metabolism may be part of high altitude adaptation, we cannot dismiss this possibility.

Second, iron deficiency can result in artificially high HbA_1c_ (via increased glycation of Hb due to elevated malondialdehyde, which is elevated during iron deficiency); this increase is significant, but small enough that it is arguably unlikely to significantly overestimate rates of diabetes [22, 29]. Nonetheless, the real possibility exists that iron deficiency increased both the probability that a participant was categorized as diabetic (by artificially increasing HbA_1c_, relative to iron replete participants), and the probability a participant was anemic. To control for this possible confounding, we included a biomarker of iron deficiency, sTfR, in all analyses, which should mitigate the risk that iron deficiency is driving our findings. We are further confident that the observed associations between diabetes and anemia are not attributable to iron deficiency because most anemic participants had non-iron deficiency anemia (8.82% of anemic Mosuo had iron deficiency; 38.13% in CHNS).

### Implications for high altitude research

The study of interactions between high altitude adaptations and chronic diseases—and diabetes in particular—is likely to become increasingly important. The Mosuo are undergoing rapid market integration, which affects diets, stress, and activity in complex ways, some of which seem to be increasing diabetes risk. Although the high altitude adaptations that are best understood involve systems that deliver oxygen to tissues, it is likely that aspects of metabolism are involved in adaptation to high altitude as well—possibly as a means to conserve the scarce resource of oxygen, or to divert it differently during exertion [6]. Genes involved in metabolism were among those identified as under selective pressure among Tibetans [6]. This raises the possibility that diabetes differs for these populations in more than just the associated anemia risk.

Further, populations native to different high-altitude regions differ in their adaptations to hypoxic stress in ways that may dramatically affect their experience of chronic disease. For example, while decoupling of hypoxia-sensing and erythropoiesis (and thus, relatively normal Hb) is part of the Tibetan suite of high altitude adaptations, elevated erythropoiesis and high Hb are part of the Andean suite. It is thus possible that this reliance on erythropoiesis to adapt to altitude leaves Andeans *more*, rather than less, vulnerable to diabetes-associated anemia than lowland natives.

### Implications for diabetes research

For the study of diabetes, further understanding of how Tibetan-type high altitude adaptations have uncoupled erythropoiesis from hypoxia may point to new therapeutic strategies to protect diabetes patients against anemia, or to mitigate the adverse effects of anemia on survival among diabetics.

